# Computation of single-cell metabolite distributions using mixture models

**DOI:** 10.1101/2020.10.07.329342

**Authors:** Mona K. Tonn, Philipp Thomas, Mauricio Barahona, Diego A. Oyarzún

**Author notes:** Corresponding author; This work was partly funded by the Human Frontier Science Program through a Young Investigator Grant (RGY0076-2015) awarded to D.O., a UKRI Future Leaders Fellowship (MR/T018429/1) awarded to P.T., and the EPSRC Centre for Mathematics of Precision Healthcare (EP/N014529/1) awarded to M.B.

## Abstract

Metabolic heterogeneity is widely recognised as the next challenge in our understanding of non-genetic variation. A growing body of evidence suggests that metabolic heterogeneity may result from the inherent stochasticity of intracellular events. However, metabolism has been traditionally viewed as a purely deterministic process, on the basis that highly abundant metabolites tend to filter out stochastic phenomena. Here we bridge this gap with a general method for prediction of metabolite distributions across single cells. By exploiting the separation of time scales between enzyme expression and enzyme kinetics, our method produces estimates for metabolite distributions without the lengthy stochastic simulations that would be typically required for large metabolic models. The metabolite distributions take the form of Gaussian mixture models that are directly computable from single-cell expression data and standard deterministic models for metabolic pathways. The proposed mixture models provide a systematic method to predict the impact of biochemical parameters on metabolite distributions. Our method lays the groundwork for identifying the molecular processes that shape metabolic heterogeneity and its functional implications in disease.

## I. INTRODUCTION

Non-genetic heterogeneity is a hallmark of cell physiology. Isogenic cells can display markedly different phenotypes as a result of the stochasticity of intracellular processes and fluctuations in environmental conditions. Gene expression variability, in particular, has received substantial attention thanks to robust experimental techniques for measuring transcripts and proteins at a single-cell resolution^1,2^. This progress has gone hand-in-hand with a large body of theoretical work on stochastic models to identify the molecular processes that affect expression heterogeneity^3–7^.

In contrast to gene expression, our understanding of stochastic phenomena in metabolism is still in its infancy. Traditionally, cellular metabolism has been regarded as a deterministic process on the basis that metabolites appear in large numbers that filter out stochastic phenomena^8^. But this view is changing rapidly thanks to a growing number of single-cell measurements of metabolites and co-factors^9–17^ that suggest that cell-to-cell metabolite variation is much more pervasive than previously thought. The functional implications of this heterogeneity are largely unknown but likely to be substantial given the roles of metabolism in many cellular processes, including growth^18^, gene regulation^19^, epigenetic control^20^ and immunity^21^. For example, metabolic heterogeneity has been linked to bacterial persistence^22,23^, a dormant phenotype characterised by a low metabolic activity, as well as antibiotic resistance^24^ and other functional effects^25^. In biotechnology applications, metabolic heterogeneity is widely recognised as a limiting factor on metabolite production with genetically engineered microbes^26–28^.

A key challenge for quantifying metabolic variability is the difficulty in measuring cellular metabolites at a single-cell resolution^29–31^. As a result, most studies use other phenotypes as a proxy for metabolic variation, e.g. enzyme expression levels^32,33^, metabolic fluxes^34^ or growth rate^35,36^. From a computational viewpoint, the key challenge is that metabolic processes operate on two timescales: a slow timescale for expression of metabolic enzymes, and a fast timescale for enzyme catalysis. Such multiscale structure results in stiff models that are infeasible to solve with standard algorithms for stochastic simulation^37^. Other strategies to accelerate stochastic simulations, such as *τ*-leaping^38^, also fail to produce accurate simulation results due to the disparity in molecule numbers between enzymes and metabolites^39^. These challenges have motivated a number of methods to optimise stochastic simulations of metabolism^40–44^. Most of these methods exploit the timescale separation to accelerate simulations at the expense of some approximation error. This progress has been accompanied by a number of theoretical results on the links between molecular processes and the shape of metabolite distributions^6,45–47^. Yet to date there are no general methods for computing metabolite distributions that can handle inherent features of metabolic pathways such as feedback regulation, complex stoichiometries, and the high number of molecular species involved.

In this paper we present a widely applicable method for approximating single-cell metabolite distributions. Our method is founded on the timescale separation between enzyme expression and enzyme catalysis, which we employ to approximate the stationary solution of the chemical master equation. The approximate solution takes the form of mixture distributions with: (i) mixture weights that can be computed from models for gene expression or single-cell expression data, and (ii) mixture components that are directly computable from deterministic pathway models. The resulting mixture model can be employed to explore the impact of biochemical parameters on metabolite variability. We illustrate the power of the method in two exemplar systems that are core building blocks of large metabolic networks. Our theory provides a quantitative basis to draw testable hypotheses on the sources of metabolite heterogeneity, which together with the ongoing efforts in single-cell metabolite measurements, will help to re-evaluate the role of metabolism as an active source of phenotypic variation.

## II. GENERAL METHOD FOR COMPUTING METABOLITE DISTRIBUTIONS

We consider metabolic pathways composed of enzymatic reactions interconnected by sharing of metabolites as substrates or products. In general, we consider models with *M* metabolites *P_i_* with *i* ∈ {1, 2, …, *M*} and *N* catalytic enzymes *E_j_* with *j* ∈ {1, 2, …, *N*}. A typical enzymatic reaction has the form

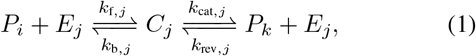

where *P_i_* and *P_k_* are metabolites, and *E_j_* and *C_j_* are the free and substrate-bound forms of the enzyme. The parameters (*k*_f, *j*_, *k*_b, *j*_) and (*k*_cat, *j*_, *k*_rev, *j*_) are positive rate constants specific to the enzyme. In contrast to traditional metabolic models, where the number of enzyme molecules is assumed constant, here we explicitly model enzyme expression and enzyme catalysis as stochastic processes. Our models also account for dilution of molecular species by cell growth and consumption of the metabolite products by downstream processes.

Though in principle one can readily write a Chemical Master Equation (CME) for the marginal distribution **P**(*P*_1_, *P*_2_, … *P_M_*) given the pathway stoichiometry, analytical solutions of the CME are tractable only in few special cases. To overcome this challenge, we propose a method for approximating metabolite distributions that can be applied in a wide range of metabolic models. We first note that using the Law of Total Probability, the marginal distribution **P**(*P*_1_, *P*_2_,…, *P_M_*) can be generally written as:

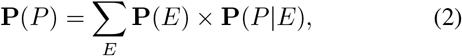

where *P* = (*P*_1_, *P*_2_, … *P_M_*) and *E* = (*E*_1_, *E*_2_, …, *E_N_*) are the vectors of metabolite and enzyme abundances, respectively. The equation in (2) describes the metabolite distribution in terms of fluctuations in gene expression, comprised in the distribution **P**(*E*), and fluctuations in reaction catalysis, described by conditional distribution **P**(*P*|*E*).

A key observation is that Eq. (2) corresponds to a mixture model with weights **P**(*E*) and mixture components **P**(*P*|*E*). To compute the mixture weights and components, we make use of the timescale separation between gene expression and metabolism. Gene expression operates on a much slower timescale than catalysis^41,45,48^, with protein half-lives typically comparable to cell doubling times and catalysis operating in the millisecond to second range. Therefore, in the fast timescale of catalysis we can write a conservation law for the total amount of each enzyme (free and bound):

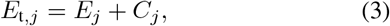

where *E*_t, *j*_ is the total number of enzymes *E_j_*. Note that since our models integrate enzyme kinetics with enzyme expression, the variables *E*_t, *j*_ follow their own, independent stochastic dynamics. It is important to note that in our approach, the conservation relation in (3) holds only in the fast timescale of catalysis. This contrasts with classic deterministic models for metabolic reactions, which typically focus on the fast catalytic timescale and assume enzymes as constant model parameters^49^.

As a result of the separation of timescales, the weights and components of the mixture in (2) can be computed separately. Specifically, the mixture weights **P**(*E*) can be obtained as solutions of a stochastic model for enzyme expression^4^, or taken from absolute single-cell measurements of enzyme expression. Such absolute measurements can be obtained from single-molecule technologies^50^, carefully calibrating fluorescence data^51,52^ or normalisation^2^. The mixture components **P**(*P*|*E*), on the other hand, can be estimated with suitable approximation techniques. For simplicity, here we choose to employ the Linear Noise Approximation (LNA), which provides a Gaussian estimate of the stationary distribution of a stochastic chemical system^53,54^. The use of the LNA is justified on the basis that metabolites tend to appear in large numbers per cell, a key condition for the LNA to produce accurate results. However, more accurate methods to compute **P**(*P*|*E*) can be used if required^60,61^. In Figure 1 we illustrate a schematic of the proposed method.

**FIG. 1.**
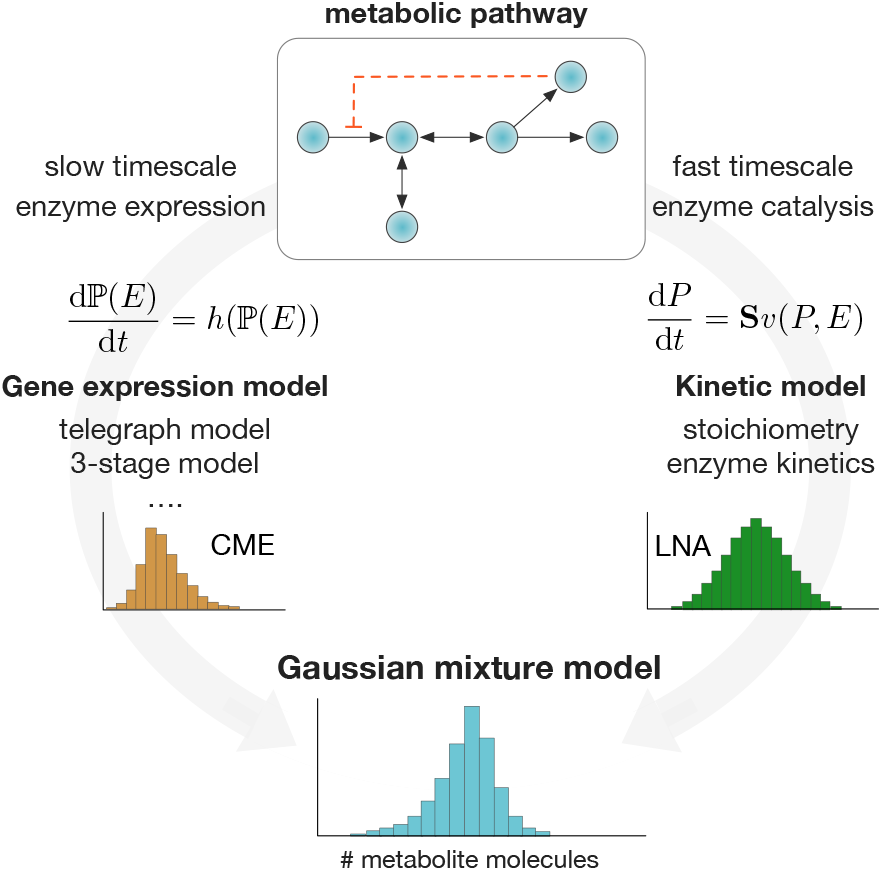
Computation of single-cell metabolite distributions with Gaussian mixture models. We exploit the separation of timescales to compute the weights and components of the mixture model in Eq. (2). Mixture weights are computed as stationary solutions to the Chemical Master Equation (CME) for a chosen model for stochastic enzyme expression. The mixture components are computed via the Linear Noise Approximation^54^ (LNA) applied to the pathway ODE model. The method produces a Gaussian mixture model for metabolite distributions that can be applied in a wide range of metabolic pathways.

We thus propose the following procedure for computing single-cell metabolite distributions:

1. Starting from the mixture model in Eq. (2), compute the enzyme distribution **P**(*E*) from a stochastic model for gene expression, either analytically (if possible) or numerically with Gillespie’s algorithm.
2. To approximate the mixture components **P**(*P*|*E*) with the LNA, compute the steady state solution 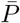 of the deterministic rate equation for each enzyme state *E*:

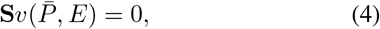

where **S** is the stoichiometric matrix and *v*(·) is the vector of deterministic reaction rates; for ease of notation we have assumed a unit cell volume, and hence the deterministic rates are equal to the propensities of the stochastic model. Note that due to the timescale separation, Eq. (4) must be solved assuming constant enzymes E, and its solution depends on the enzyme abundance, i.e. 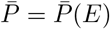.
3. For each enzyme state *E*, compute the solution to the Lyapunov equation^54^:

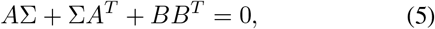

where *A* is the Jacobian of (4) evaluated at the steady state and *BB^T^* = **S**diag {*v*} **S**^*T*^. Note that, as in (4), the solution of the Lyapunov equation depends on the enzyme state, i.e. Σ = Σ(*E*).
4. Following the LNA, approximate the mixture components **P**(*P*|*E*) as a multivariate Gaussian distribution with mean 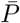 and covariance matrix Σ.
5. Combine the weights **P**(*E*) and Gaussian components **P**(*P*|*E*) through the mixture model in (2).

In the next sections we illustrate the effectiveness of our method in two exemplar systems.

## III. REVERSIBLE MICHAELIS-MENTEN REACTION

We first consider a stochastic model that integrates a reversible Michaelis-Menten reaction with a standard model for enzyme expression. As shown in Figure 2A, the Michaelis-Menten mechanism includes reversible binding of four species: a metabolic substrate *S*, a free enzyme *E*, a substrate-enzyme complex *C* and a metabolic product *P*. To model enzyme expression, we use the well-known two-stage scheme for transcription and translation^55,56^ (Figure 2A). The complete set of reactions is:

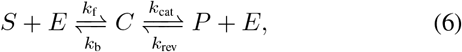

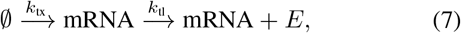

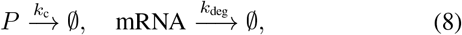

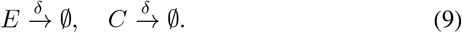

**FIG. 2.**
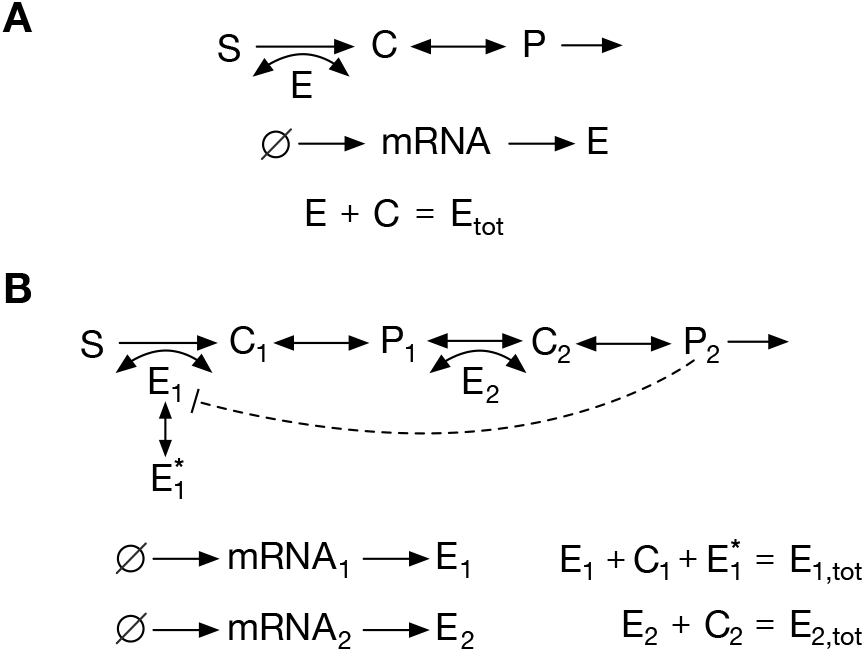
Exemplar metabolic systems. (**A**) Reversible Michaelis-Menten reaction; the full set of reactions are shown in Eq. (6)–(9). The model accounts for reversible catalysis of a substrate *S* into a product *P*. (**B**) Two-step pathway with noncompetitive end-product inhibition; the reactions are shown in Eq. (18)–(25). The product (*P*_2_) sequesters enzyme *E*_1_ into an inactive form 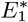, thereby reducing the rate of the first reaction. In both examples we assume a constant substrate *S* and linear dilution of all chemical species. Enzymes are assumed to follow the two-stage model for gene expression^56^, which includes species for the enzymatic mRNA and protein.

The reactions in (6) correspond to a reversible Michaelis-Menten reaction as in (1), while reactions in (7) are the two-stage model for gene expression. We include four additional first-order reactions (8)–(9) to model consumption of the metabolite product with rate constant *k_c_*, mRNA degradation with rate constant *k*_deg_, and dilution of all model species with rate constant *δ*. In what follows we assume that the substrate *S* remains strictly constant, for example to model cases in which the substrate represents an extracellular carbon source that evolves in much slower timescale than cell doubling times.

Since on the fast timescale of the catalytic reaction, the total number of enzymes can be assumed in quasi-stationary state^6,49^, we have that

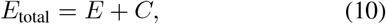

and therefore the general mixture model in (2) can be written as:

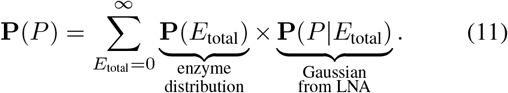

The mixture weights **P**(*E*_total_) can be computed from the stochastic model for gene expression in (7). Under the standard assumption that mRNAs are degraded much faster than proteins^4^, the stationary solution of the two-stage model can be approximated by a negative binomial distribution^56^:

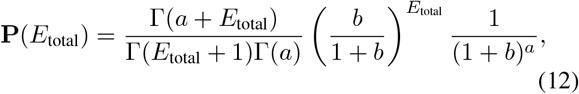

where Γ is the Gamma function and the parameters are defined as the burst frequency *a* = *k*_tx_/*δ* and burst size *b* = *k*_tl_ /*k*_deg_.

To compute the mixture components **P**(*P*|*E*_total_) with the LNA, we write the full system of deterministic rate equations (see (35) in Methods) for the three species *E*, *C* and *P*. Note that in this case, we can further reduce the rate equations by (i) using the conservation law in (10), and (ii) assuming that the binding and unbinding reactions between *S* and *E* reach equilibrium faster than the product *P*, a condition that generally holds in metabolic reactions. After algebraic manipulations, the reduced ODE can be written as:

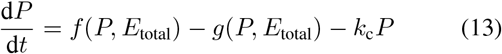

where

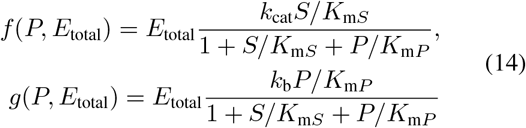

and the parameters are *K*_m*S*_ = (*k*_b_ + *k*_cat_)/*k*_f_ and *K*_m*P*_ = (*k*_b_ + *k*_cat_)/*k*_rev_.

The mean of each mixture component is simply given by the steady state solution of (13), which we denote as 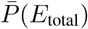. For a given enzyme abundance *E*_total_, the variance ∑(*E*_total_) of each Gaussian component is given by the solution to the Lyapunov equation in (5):

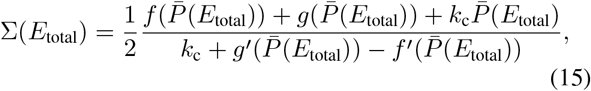

where *f*′ and *g*′ are first-order derivatives. Combining the negative binomial in (12) with the Gaussian components, we can rewrite Eq. (11) to get a Gaussian mixture model for the metabolite:

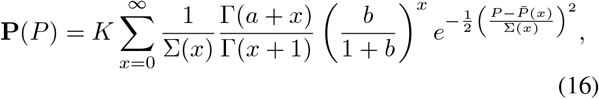

where both 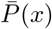 and Σ(*x*) must be computed for each value of *x* = *E*_total_ in the summation. The normalization constant in (16) is

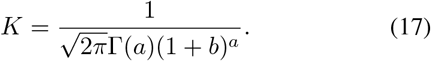

In Figure 3 we plot the mixture model (16) for realistic parameter values and compare this approximation with distributions computed from long runs of Gillespie simulations of the whole set of reactions (6)–(9). The results indicate that the mixture model provides an excellent approximation of the metabolite distribution. In the next section we test our methodology in a more complex pathway with feedback regulation.

**FIG. 3.**
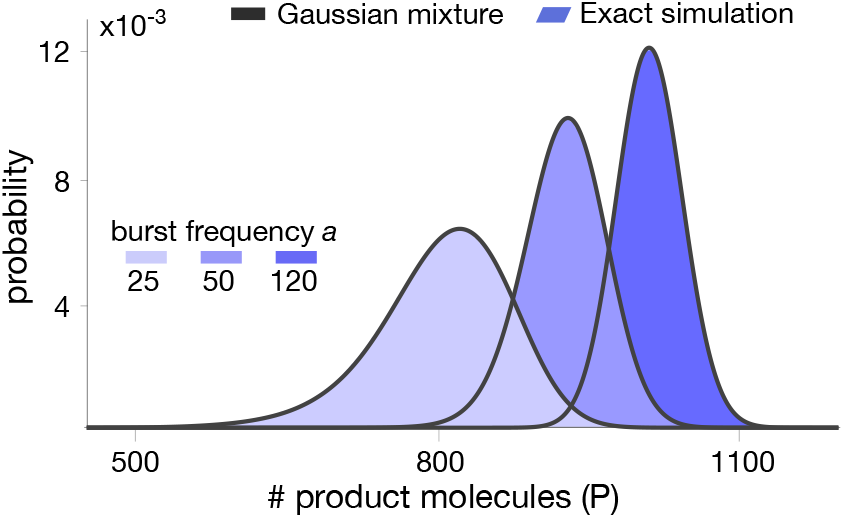
Stationary product distribution of a Michaelis-Menten reaction. The proposed mixture model in (16) provides an excellent approximation for the metabolite distribution obtained with Gillespie’s algorithm^37^. Distributions were computed for varying values of the bursting parameter *a*. Note that the resulting distributions are almost identical to those predicted in our earlier work using a Poisson mixture^6^, since we have deliberately chosen parameters to produce similar distribution in both cases. All parameter values can be found in Table I in the Methods.

**TABLE I.** Parameter values for simulations in Figure 3.

| Figure 3 |  |  |  |
| --- | --- | --- | --- |
| $\delta$ | $0.00025 \text{ s}^{-1}$ | $k_b$ | $1000 \text{ s}^{-1}$ |
| $a$ | $\{25, 50, 120\}$ | $k_{\text{cat}}$ | $3.6 \text{ s}^{-1}$ |
| $b$ | 1 | $k_{\text{rev}}$ | $0.01 \text{ s}^{-1}$ |
| $S$ | 3000 molecules | $k_c$ | $0.02 \text{ s}^{-1}$ |
| $k_f$ | $1 \times S \text{ s}^{-1}$ | | |

## IV. PATHWAY WITH END-PRODUCT INHIBITION

A common regulatory motif in metabolism is end-product inhibition, in which a pathway enzyme can bind to its own substrate as well as the pathway product (see Figure 2B). The product thus sequesters enzyme molecules, which reduces the number of free enzymes available for catalysis and slows done the reaction rate. To examine the accuracy of our method in this setting, we study a fully stochastic model for a two-step pathway with noncompetitive end-product inhibition:

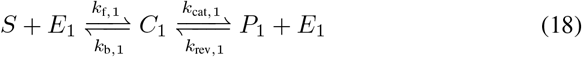

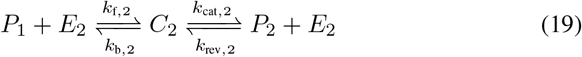

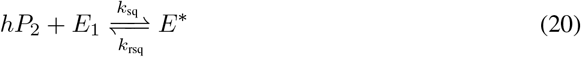

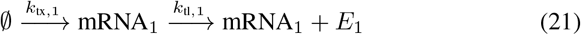

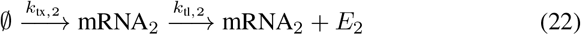

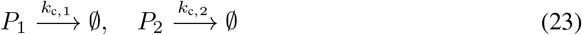

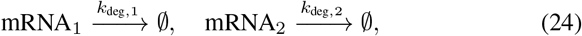

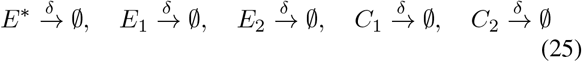

The two reactions in (18) and (19) are reversible Michaelis-Menten kinetics, sharing the intermediate metabolite *P*_1_ as a product and substrate, respectively. The end-product inhibition in (20) consists of reversible binding between *h* molecules of *P*_2_ and the first enzyme *E*_1_ into a catalytically-inactive complex *E*\*. The remaining model reactions in (21)–(25) are analogous to the previous example in Section III: reactions in (21)–(22) describe the two-stage model for expression of both enzymes, and with reactions (23)–(25) we model first-order mRNA degradation, product consumption, and dilution by cell growth. For simplicity we also assume that both enzymes are independently expressed, but in general our method can also account for cases in which enzymes are co-expressed or co-regulated^57^. The resulting model has two distinct pools of enzymes, which remain constant over the timescale of catalysis:

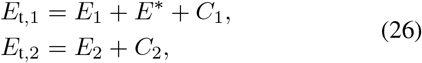

and therefore the mixture model in (2) becomes

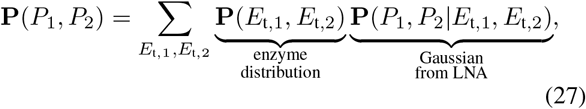

where the summation goes through all (*E*_t,1_, *E*_t, 2_) pairs. Since both enzymes are expressed independently, the enzyme distribution is the product of two negative binomials **P**(*E*_t, 1_, *E*_t, 2_) = **P**(*E*_t, 1_) × **P**(*E*_t, 2_), each one analogous to the distribution in (12).

To compute the mixture components with the LNA, we use the rate equations for the reactions in (18)–(23); the full set of ODEs is listed in Eq. (36) in the Methods. As in the first example, by employing the conservation laws in (26) and assuming rapid equilibrium of the complexes *C*_1_ and *C*_2_, the deterministic model can be further simplified to a 2-dimensional ODE:

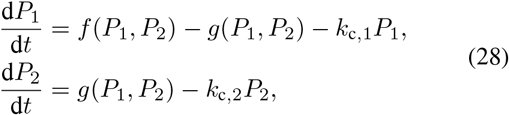

where for ease of notation we have omitted the dependency on *E*_t, 1_ and *E*_t, 2_. The nonlinear functions in (28) are

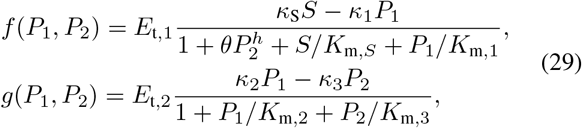

where *θ* = *k*_sq_/*k*_rsq_ is the product-enzyme binding constant and the remaining parameters are defined as *κ*_S_ = *k*_cat, 1_*k*_f,1_/(*k*_b,1_ + *k*_cat, 1_), *κ*_1_ = *k*_b,1_*k*_rev,1_/(*k*_b,1_ + *k*_cat,1_), *κ*_2_ = *k*_cat,2_*k*_f,2_/(*k*_b,2_ + *k*_cat,2_), *κ*_3_ = *k*_b,2_*k*_rev,2_/(*k*_b,2_ + *k*_cat,2_), *K*_m, *S*_ = *k*_cat,1_/ *κ*_S_, *K*_m,1_ = *k*_b,1_/*κ*_1_, *K*_m,2_ = *κ*_cat,2_/ *κ*_2_ and *K*_m,3_ = *k*_b,2_/*κ*_3_.

As in the previous example, the ODEs in (28) correspond to the full model (36) rewritten in terms of both metabolites assuming that the enzyme-substrate reactions reach equilibrium in a faster timescale than catalysis. This reduced model can be readily employed to obtain approximations for the mixture components with the LNA. If we denote as 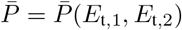 the steady state solution of (28), we can write the Lyapunov equation as *A*Σ + Σ*A^T^* + *BB^T^* = 0 with *A* and *BB^T^* given by

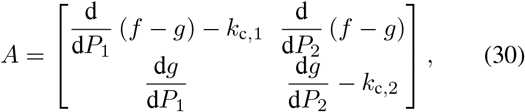

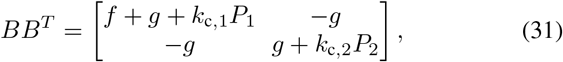

where *f*(·), *g*(·), and their derivatives are evaluated at the steady state solution 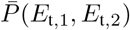. The Gaussian components of the mixture model are then

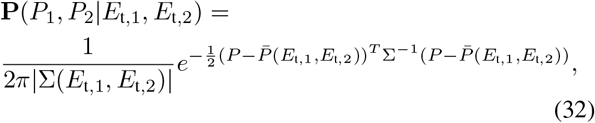

where *P* = (*P*_1_, *P*_2_)^*T*^ and |·| is the matrix determinant. After combining the joint distribution of enzymes and the components into Eq. (27), we get a Gaussian mixture model for the joint marginal distribution of both metabolites:

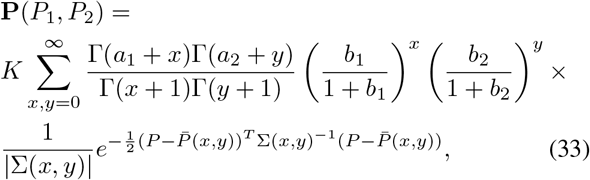

where 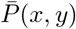 and Σ(*x*, *y*) need to computed numerically for each pair (*x*, *y*) = (*E*_t, 1_, *E*_t, 2_) in the summation. The burst frequencies *a_i_* = *k*_tx, *i*_/*δ* and burst sizes *b_i_* = *k*_tl, *i*_/*k*_deg, *i*_ are specific to each enzyme, and the normalisation constant is given by

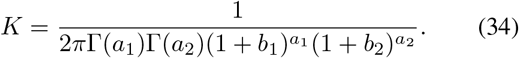

To test the quality of the approximation, we numerically computed the mixture model in (33) for various combinations of parameter values, shown in Figure 4. We observe that the mixture model offers an excellent approximation as compared to exact Gillespie simulations of the full model (18)–(25). We note that in this case, the full stochastic model has seven species and three different timescales, and therefore the runtime of Gillespie simulations are extremely long, in the order of several hours per run.

**FIG. 4.**
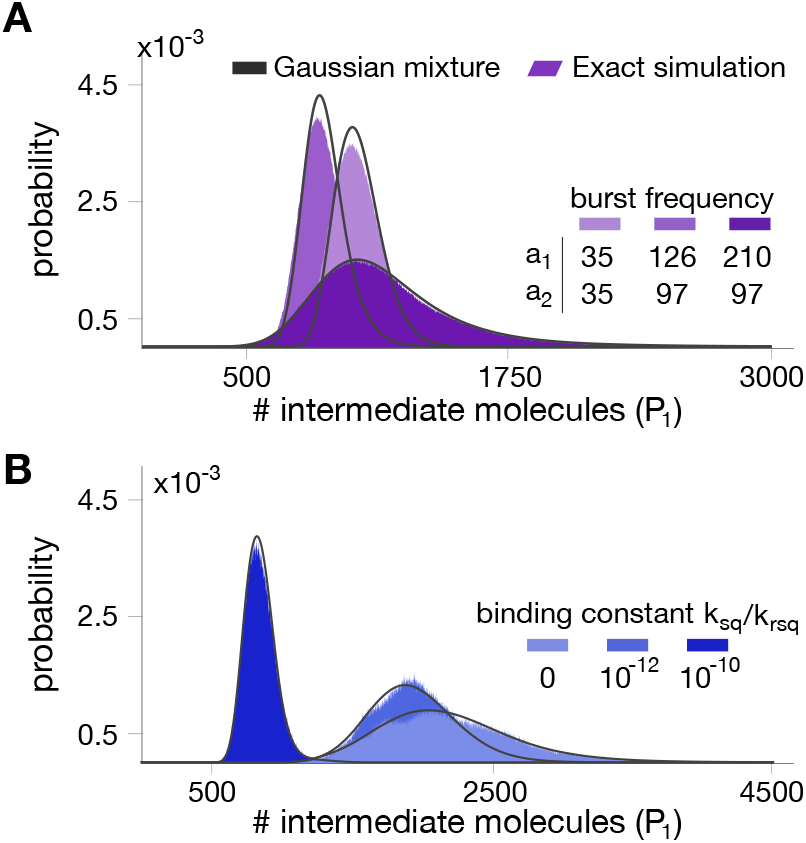
Stationary distributions for the intermediate metabolite in a two-step pathway with end-product inhibition. The panels show the distribution of intermediate metabolite *P*_1_ for different combinations of parameter values. **(A)** Impact of enzyme bursting frequency *a*_1_ and *a*_2_. **(B)** Impact of binding constant between the first enzyme and the end-product. All parameter values can be found in Table II in the Methods.

**TABLE II.** Parameter values for simulations in Figure 4.

| Figure 4 |  |  |  | Figure 4A |  | Figure 4B |  |
| --- | --- | --- | --- | --- | --- | --- | --- |
| $\delta$ | $0.00025 \text{ s}^{-1}$ | $k_{\text{rev},1}$ | $0.0001 \text{ s}^{-1}$ | $a_1$ | $\{35, 126, 210\}$ | $a_1$ | 80 |
| $k_{\text{deg},1}$ | $0.2 \text{ s}^{-1}$ | $k_{c,1}$ | $0.00025 \text{ s}^{-1}$ | $a_2$ | $\{35, 97, 97\}$ | $a_2$ | 80 |
| $k_{\text{deg},2}$ | $0.2 \text{ s}^{-1}$ | $k_{f,2}$ | $1.5 \text{ s}^{-1}$ | $b_1$ | 1 | $b_1$ | 1 |
| $S$ | 3000 molecules | $k_{b,2}$ | $15000 \text{ s}^{-1}$ | $b_2$ | 1 | $b_2$ | 1 |
| $k_{f,1}$ | $20 \times S \text{ s}^{-1}$ | $k_{\text{cat},2}$ | $150 \text{ s}^{-1}$ | $k_{\text{sq}}$ | $10^{-10} \text{ s}^{-1}$ | $k_{\text{sq}}$ | $\{0, 10^{-10}, 10^{-12}\} \text{ s}^{-1}$ |
| $k_{b,1}$ | $15000 \text{ s}^{-1}$ | $k_{\text{rev},2}$ | $0.001 \text{ s}^{-1}$ | $k_{\text{rsq}}$ | $1 \text{ s}^{-1}$ | $k_{\text{rsq}}$ | $1 \text{ s}^{-1}$ |
| $k_{\text{cat},1}$ | $22.5 \text{ s}^{-1}$ | $k_{c,2}$ | $0.15 \text{ s}^{-1}$ | $h$ | 3 | $h$ | 3 |

To further illustrate the utility of our method, we employed the mixture model to study the impact of parameter perturbations on the metabolite distributions. Without an analytical solution, such a study would require the computation of long Gillespie simulations for each combination of parameter values, which quickly become infeasible due to the long simulation time. In contrast, the mixture model provides a systematic way to rapidly evaluate the influence of model parameters on metabolite distributions. In Figure 5A we show summary statistics of the marginal **P**(*P*_1_) for various combinations of average enzyme expression levels. The results suggest that expression levels can have a strong impact on the mean and coefficient of variation of the intermediate metabolite. Moreover, in Figure 5B we plot the distribution **P**(*P*_1_, *P*_2_) for combinations of bursting parameters. The results show that uncorrelated enzyme fluctuations can result in correlated metabolite distributions due to the coupling introduced by the pathway^45^.

**FIG. 5.**
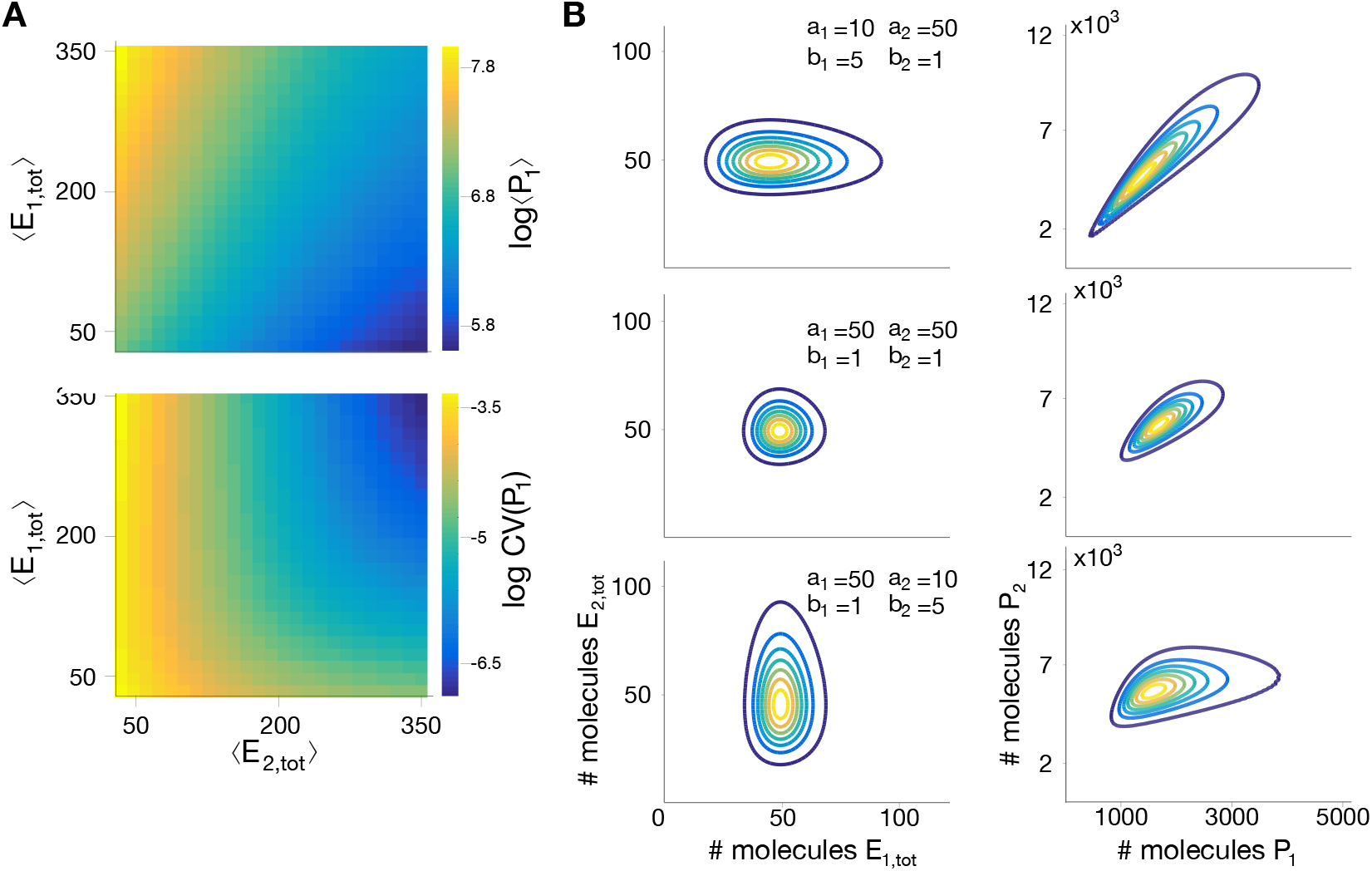
Impact of enzyme expression on metabolite distributions. **(A)** We compute the mean and coefficient of variation (CV) of the intermediate metabolite *P*_1_ in model (18)–(25), for a wide range of mean enzyme expression levels. **(B)** Enzyme expression parameters shape the metabolite distribution; We computed the joint metabolite distribution **P**(*P*_1_, *P*_2_) for three combinations of enzyme bursting parameters, chosen to give the same mean expression, and assuming both enzymes are expressed independently. Shown are contour plots of the bivariate distributions of enzymes (left) and metabolites (right). The results suggest that metabolite correlations emerge even when enzymes are uncorrelated, as reported previously in the literature^45^. All parameter values can be found in Table III in the Methods.

**TABLE III.** Parameter values for simulations in Figure 5.

| Figure 5A |  | Figure 5B |  |
| --- | --- | --- | --- |
| $a_1$ | $[10, 100]$ | $a_1$ | $\{10, 50, 50\}$ |
| $a_2$ | $[10, 100]$ | $a_2$ | $\{50, 50, 10\}$ |
| $b_1$ | 1 | $b_1$ | $\{5, 1, 1\}$ |
| $b_2$ | 1 | $b_2$ | $\{1, 1, 5\}$ |
| $k_{\text{sq}}$ | $10^{-10} \text{ s}^{-1}$ | $k_{\text{sq}}$ | $0 \text{ s}^{-1}$ |
| $k_{\text{rsq}}$ | $1 \text{ s}^{-1}$ | $k_{\text{rsq}}$ | $1 \text{ s}^{-1}$ |
| $h$ | 3 | $h$ | 3 |

## V. DISCUSSION

Cellular metabolism has traditionally been assumed to follow deterministic dynamics. This paradigm results largely from the observation that cellular metabolites are highly abundant. However, recent data shows that single-cell metabolite distributions can display substantial heterogeneity in their abundance across single cells^9–17^. It has also been shown that expression of metabolic genes is as variable as any other component of the proteome^2^, and thus in principle it is plausible that such enzyme fluctuations propagate to metabolites. These observations have begun to challenge the paradigm of metabolism being a deterministic process, suggesting that metabolite fluctuations may play a role in non-genetic heterogeneity.

Here we described a new computational tool to predict the statistics of metabolite fluctuations in conjunction with gene expression. The method is based on a timescale separation argument and leads to a Gaussian mixture model for the stationary distribution of cellular metabolites. Computing distributions from this approximate model is substantially faster than through stochastic simulations, as these can be extremely slow due to the multiple timescales of metabolic pathways. Our technique can therefore be employed to efficiently explore the parameter space and predict the shape of metabolite distributions in different conditions. In earlier work we showed that the product of a single metabolic reaction can be accurately described by a Poisson mixture model^6^. Such approximation allowed the discovery of previously unknown regimes for metabolite distributions, including heavily tailed distributions and various types of bimodality and multimodality. The Poisson approximation, however, is bespoke to single reactions and not valid for more complex systems. In contrast, the Gaussian mixture model discussed here can be applied to multiple kinetic mechanisms, more complex stoichiometries, as well as post-translational regulation.

An advantage of our approach is that the mixture weights can be computed offline from stochastic models for gene expression or single-cell expression data. The model is flexible in that it can readily accommodate gene expression models of various complexity. For the sake of illustration, in our examples we used the simple two-stage model for gene expression, but other models including gene regulation can also be employed^7^. Particularly relevant models are those that account for enzyme co-regulation, a widespread feature of bacterial operons^57^, which translates into correlations between expression of different pathway enzymes and the resulting metabolite abundances. A limitation of our method is that in many cases analytic solutions of the CME are not known, particularly for large models with multiple interacting genes. In such cases, the mixture weights **P**(*E*) can be approximated through stochastic simulations^37^ albeit at the expense of increased computational costs. Most recently, progress in stochastic simulation of genome-scale metabolic networks^58^ can offer an alternative route for studying fluctuations in large metabolic models.

The effectiveness of our method relies on two conditions: the separation of timescales between enzyme expression and enzyme catalysis, and the ability of the LNA to approximate the mixture components accurately. The first condition is satisfied by the vast majority of enzymes because their kinetics operate in regimes that are orders of magnitude faster than gene expression^57^. However, the timescale separation can fail if the metabolic substrate S, typically a carbon source, cannot be assumed to be constant, a suitable assumption in the typical case of abundant nutrient sources with low fluctuations. Our theory would need to be extended in cases when nutrient sources become another source of variability, e.g. under fluctuations dictated by the environment^7^. The second condition breaks down when the LNA fails to provide good estimates of the mixture components^59,60^. As explained in Section II, here we have deliberately chosen to employ the LNA because it provides a simple and rapid method to compute the mixture components, **P**(*P*|*E*), for a broad range of metabolic pathways. Yet in cases where its assumptions do not hold, e.g. low abundance of metabolites, the LNA step in our method can be replaced by more accurate approximations. Such alternative methods include, for example, the conditional system size expansion including terms beyond the LNA, maximum entropy reconstructions using the method of conditional moments, or the finite state projection algorithm^60,61^, all of which can be readily incorporated into our mixture model strategy. These methods rely on different assumptions and their approximation quality will vary depending on the specific model parameters; in some cases, estimates for their approximation errors can be obtained with suitable methods, as discussed in a recent review on this topic^62^.

Although our method can account for a large class of metabolic models and post-translational regulation mechanisms, there are a number of promising extensions that would broaden its utility in light of recent experimental advances. First, here we have only considered stationary distributions of metabolites, and a number of experiments have revealed cases in which metabolic heterogeneity emerges during dynamic nutrient shifts^32,33,63^. Extensions of our method to time-dependent metabolite distributions require the computation of the time-dependent solution of the CME for the enzyme expression model^56,64^. As long as the dynamics of gene expression is slow enough to preserve the time scale separation, the computation of the mixture components with the LNA or other methods remains unchanged.

Another promising extension is the inclusion of transcriptional feedback regulation, a topic that has received substantial attention in the literature^19,65–67^. In these systems, some pathway metabolites can bind to transcription factors (TF) that control enzyme expression in the same pathway. Such regulation can be included by using the conditional LNA method^68^ at the expense of not being able to compute the mixture weights offline anymore. Specifically, this extension would model mixture weights through more elaborate enzyme expression models in which the metabolite-TF interactions are replaced by their conditional averages, leading to an effective feedback model that requires specialised solution methods^69^. A particularly promising application of such extended analysis is in synthetic biology, where there is a growing interest in the interplay between stochastic fluctuations and experimentally tunable parameters of molecular circuits^70,71^. In particular, the use of metabolite-responsive feedback can improve robustness of strains engineered for the production of high-value metabolites^72,73^. Early results in this area^47^ suggest complex dependencies between metabolite fluctuations and the tunable parameters of the feedback control system. Such analyses were purely based on lumped models for metabolite-TF binding, and hence a more detailed theory could reveal novel design strategies to mitigate metabolite heterogeneity in production strains.

A number of works have sought to find links between fluctuations across layers of cellular organisation, such as gene expression, metabolism and cell growth^32,33,35,63,74^. But since measurement of metabolites in single cells remains technically challenging, there is pressing need for computational methods to predict fluctuations in cellular metabolites. Our proposed method provides a systematic approach for such task, paving the way for the generation of hypotheses on the molecular sources of metabolic heterogeneity.

## VI. METHODS

### A. Model simulation

Stochastic simulations were computed with Gillespie’s algorithm over long simulation times (several hours) corresponding to thousands of cell cycles. The ODE models and Lyapunov equations were solved in Matlab. In all examples, the negative binomial distribution for gene expression in (12) was computed with its continuum approximation (Gamma distribution).

### B. Deterministic rate equations

#### a. Reversible Michaelis Menten

The full set of rate equations for the reversible reaction in (6)–(9) is:

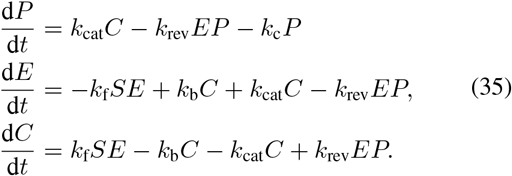

To further reduce the above system of ODEs to Eq. (13) in the main text, we can substitute the conservation relation in Eq. (10), i.e. *C* = *E*_tota1_ – *E*, and use the fact that the substrate-enzyme complex (*C*) typically equilibrates much faster than the product *P*, which means that d*C*/d*t* ≈ 0 in the timescale of catalysis.

#### b. End-product inhibition

The full set of rate equations for the reactions in (18)–(23) is:

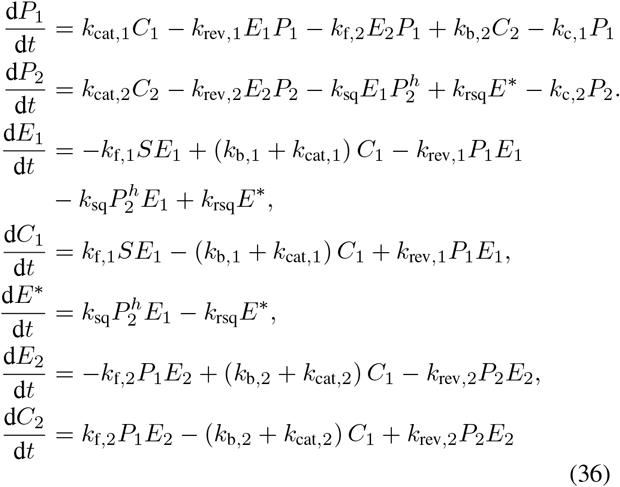

As in the previous example, we can use the rapid equilibrium assumption and the conservation relations in (26), i.e. *E*_t, 1_ = *E*_1_ + *E*\* + *C*_1_ and *E*_t, 2_ = *E*_2_ + *C*_2_, to simplify the 7-dimensional ODE in (36) to the 2-dimensional system in (28) of the main text.

## Notes

### Competing Interest Statement

The authors have declared no competing interest.

